# Error modelling of SNVs and indels in deep targeted cell-free DNA for sensitive detection of tumor burden in colorectal cancer

**DOI:** 10.64898/2026.05.29.728727

**Authors:** Mathilde Hartvig Diekema, Mads Heilskov Rasmussen, Simon Opstrup Drue, Amanda Frydendahl, Claus Lindbjerg Andersen, Jakob Skou Pedersen

## Abstract

Circulating tumor DNA (ctDNA) is a promising biomarker for cancer detection, but low tumor burden makes it difficult to distinguish true signal from background noise.

To aggregate and better evaluate weak mutational signals, we propose PyDREAMS, which incorporates both single-nucleotide variants (SNVs) and insertion and deletions (indels) for ctDNA detection and quantification. To distinguish signal from noise, a neural network background error model is learned from healthy controls. It captures the joint effects of cell-free DNA (cfDNA)-specific lesions and sequencing errors, accounting for both genomic context and read-level features. Finally, a statistical test is used to evaluate the presence of mutational signals. We evaluate the method in a tumor-informed setting, using cohorts of colorectal cancer samples with deep targeted plasma cfDNA sequencing across 12 cancer driver genes.

We trained PyDREAMS on 46 healthy controls, with feature analysis revealing that both SNV and indel error rates were lowest at mononucleosomal fragment lengths, suggesting that nucleosomes protect cfDNA and reduce lesion accumulation during circulation and sample handling. In the validation cohort, combining SNVs with indels improved detection, with indels contributing approximately 1.5-fold more evidence per mutation than SNVs. On a test cohort of 209 stage I–III colorectal cancer (CRC) patients and 24 healthy controls, PyDREAMS outperformed a Shearwater-based caller (AUC 0.917 vs 0.909). In stage III post-operative (post-OP) samples (*n* = 26), where ctDNA was expected only in non-cured patients, PyDREAMS detected ctDNA in 5 patients, including 3/9 with later recurrence, while Shearwater detected none.

Together, these results show that PyDREAMS improves evaluation of ultra-low-frequency tumor signals through unified read-level modelling of SNV and indel background error.

## 1 Introduction

Cell-free DNA (cfDNA) consists of short DNA fragments that originate from cells undergoing apoptosis or necrosis. These fragments are released into the bloodstream and have a short half-life [20, 28, 34]. Although most of cfDNA originate from hematopoietic cells, however, in patients with cancer, a minor fraction in cancer patients can be traced to tumor cells, referred to as circulating tumor DNA (ctDNA) [1, 16]. In early-stage cancers or in patients with small tumor burdens, ctDNA often comprises less than 0.1% of the total pool of cfDNA [20].

In principle, next generation sequencing (NGS) enables the detection of ctDNA, which is characterized by low-frequency somatic mutations, copy-number alterations, methylation changes, or fragmentation patterns [17, 23]. Sequencing of cfDNA from blood samples may provide a non-invasive opportunity for real-time cancer detection and disease monitoring [3, 15]. Despite this promise, ctDNA detection remains challenging in many clinical contexts, especially minimal residual disease (MRD) after curative-intent surgery and early-stage disease when the tumor burden is low [5, 34, 35].

The clinical value of ctDNA as a biomarker depends on accurately distinguishing true tumor signal from background noise. At low ctDNA fractions, genuine tumor-derived signals can be obscured by technical noise from sequencing errors [21].

For somatic mutations, confidence in variant calls increases with effective molecular depth and resulting number of molecules supporting the alternate allele [25, 20, 12]. To achieve this, many current ctDNA applications rely on deep targeted sequencing, which captures rare variant alleles by focusing on a predefined panel of genomic regions at very high depth [33, 11, 9, 19, 20]. Furthermore, ultra-deep sequencing often employs unique molecular identifiers (UMIs), allowing reads from the same original molecule to be collapsed into a consensus read that corrects most random PCR and sequencing errors [12, 20]. However, DNA damage introduced before amplification can persist after UMI consensus calling and may still generate systematic artifacts.

Two broad approaches have emerged for distinguishing true low-frequency variants from residual background noise in deep targeted cfDNA sequencing. The first relies on position-specific background tables, estimating per-locus error rates from a panel of normal (PON) and flagging sites where observed variant counts exceed these expectations. The second learns per-read error probabilities from cfDNA-specific features, exploiting that fragment structure, sequencing-cycle effects, and local sequence context may directly shape technical error rates and therefore carry information relevant for background modelling [17, 28]. Christensen et al. [6] developed deep read-level modelling of sequencing-errors (DREAMS) as a framework of the second type: a ctDNA detection approach based on a neural network that predicts mismatch and match probabilities from read-level and local sequence-context features. Training is performed on healthy controls or on patient samples after excluding ctDNA-positive positions, enabling error patterns to be learned from sites assumed to match the reference or from unmutated regions of cfDNA. By incorporating features such as read position, fragment length, UMI group size, GC content, and local trinucleotide context, DREAMS down-weights likely technical artifacts and increases confidence in non-reference observations with low predicted error rates [6].

Here, we extend DREAMS from single-nucleotide variant (SNV)-only modelling to a uni-fied framework that also incorporates insertion and deletions (indels). In tumor-informed panel sequencing, the patient-specific catalog is constrained to variants observed in the primary tumor within a fixed panel, so every informative locus contributes meaningfully to the sample-level signal. In our colorectal cancer (CRC) cohort, more than half of patients carried at least one indel within the targeted panel, meaning an SNV-only framework discards a substantial fraction of the available evidence. Moreover, indels have qualitatively different error characteristics from SNVs: they are more sensitive to alignment ambiguity, low-complexity sequence, and read-end artefacts [18, 7], and cannot simply be captured by SNV-style noise assumptions. Extending DREAMS to indels therefore requires careful handling of indel representation and separate modelling of distinct read-level error processes. Here, Python deep read-level modelling of sequencing-errors (PyDREAMS) extends DREAMS to unified read-level error modeling for both SNVs and indels, with separate neural networks for each variant class.

To evaluate PyDREAMS, we benchmarked it against ultra-deep mutation integrated sequencing (UMIseq) [10], an ultra-deep, tumor-informed, fixed-panel ctDNA assay for CRC. UMIseq combines UMI consensus reads with a PON to model locus-specific error rates using a Shearwater-based beta-binomial framework and aggregates evidence across a patient-specific variant catalog into a sample-level score. Unlike PyDREAMS, UMIseq relies on per-position error tables rather than learning per-read error probabilities from cfDNA-specific features. Evaluating both methods on the same cohorts and sequencing data as in the UMIseq study [10] allows assessment of whether a transferable, feature-based read-level error model can match detection performance while providing a unified framework for both SNVs and indels.

## 2 Results

### 2.1 Study overview

The overall workflow is shown in Figure 1. cfDNA samples from 46 healthy controls in the training cohort were sequenced and processed as described by Frydendahl et al. [10], using a custom hybrid-capture panel covering 15,465 base pairs across 42 exonic regions in 12 genes commonly altered in CRC (Figure 1a). The resulting UMI-collapsed data had a median depth of 9,086 (interquartile range (IQR) 5,464).

**Figure 1.**
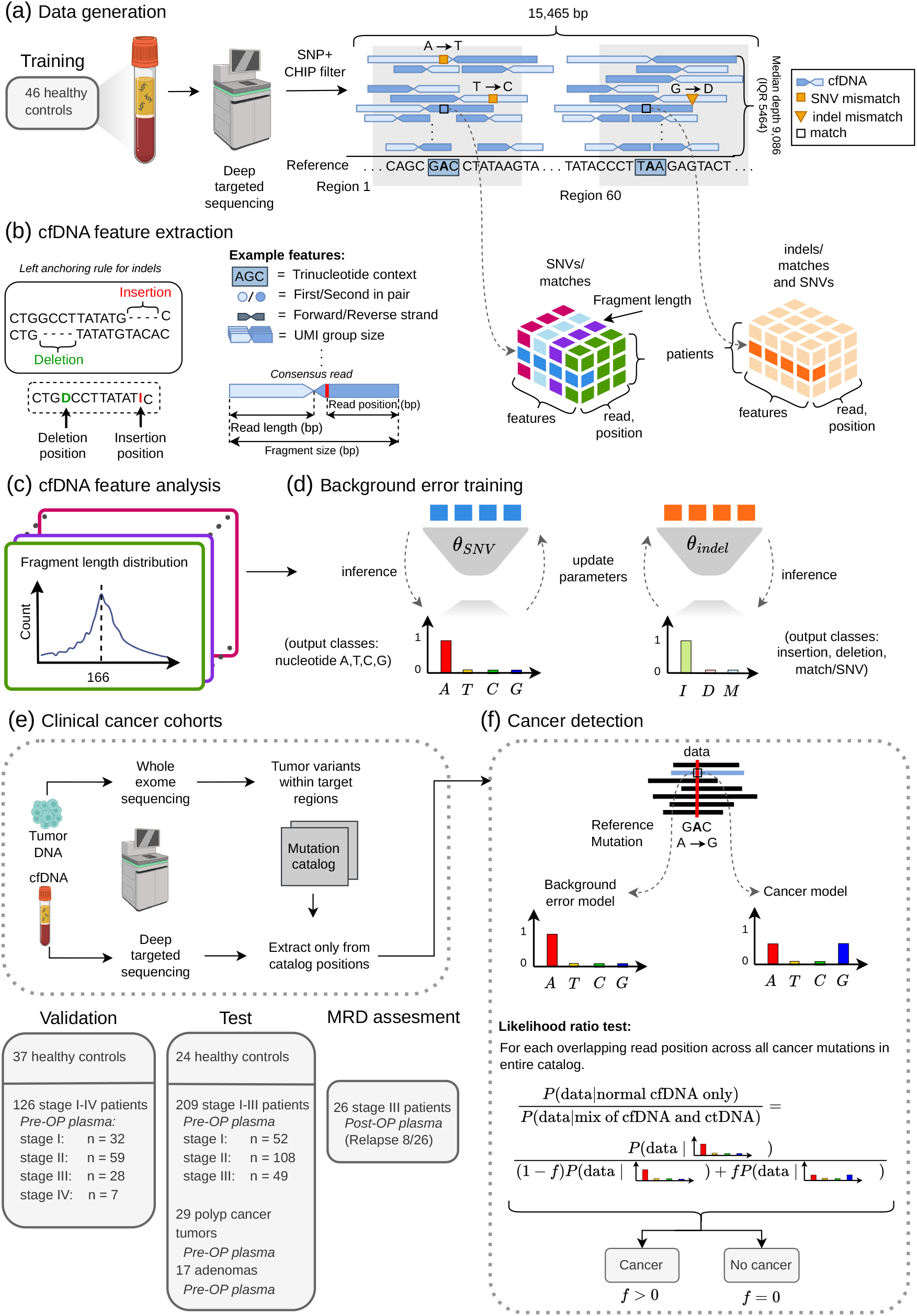
Study overview. **(a) Data generation**. Reads were classified as SNV mismatches, indel mismatches, or matches to the reference genome. A SNP and CHIP filter was applied to reduce germline and clonal hematopoietic signal. **(b) cfDNA feature extraction**. Indel candidates were assigned genomic coordinates by a left-anchoring rule: insertions were anchored to the reference base immediately 5^*′*^ upstream of the inserted sequence, and deletions to the first reference base missing from the read. Local sequence context and read-level features were extracted for each position (Figure 2a) and organized into three-dimensional tensors (patients × features × read positions) separately for SNVs and indels. **(c) cfDNA feature analysis**. Fragment length distribution shown as an example of the structured variation in cfDNA error profiles. **(d) Background error training**. Iterative training of the SNV and indel background error models (*θ*_SNV_ and *θ*_indel_) on healthy-control data. **(e) Clinical cancer cohorts**. Patient-specific mutational catalogs were constructed from WES of primary tumor biopsies and matched PBMCs, and cfDNA plasma was sequenced using the same hybrid-capture panel with reads filtered to catalog positions. Cohort composition of the validation, test, and MRD assessment subsets is shown. **(f) Cancer detection**. Likelihood ratio test comparing a background-only model to a mixture model with tumor-derived ctDNA fraction *f*, applied across all read positions overlapping the mutation catalog. Samples with *f >* 0 are classified as ctDNA-positive; samples with *f* = 0 as ctDNA-negative.

Training data for the null model of background sequencing errors were generated by extracting local sequence context and read-level features from all mismatches to the reference genome in the training cohort, together with a random subsample of matches, using the same approach as Christensen et al. [6] (Figure 1b). As a data preprocessing step, a two-stage background mask was subsequently applied to reduce residual germline signal and identify recurrent technical noise. The same step was also done for test samples (Methods, Section 5.2).

Using the resulting datasets, we performed an exploratory analysis of the extracted features to identify and characterize structured variation in cfDNA error profiles (Figure 1c and 2).

Two separate neural networks were trained on the healthy-control data to learn background error models *θ*_SNV_ and *θ*_indel_ (Figure 1d). The SNV network predicted per-read error probabilities over the four nucleotides (A, T, C, G), while the indel network predicted class probabilities for insertions (I), deletions (D), and matches or mismatches (M). Parameters were optimized by minimizing the discrepancy between predicted and observed error profiles in the training cohort.

For the clinical cohorts, patient-specific mutational catalogs were constructed by performing whole-exome sequencing (WES) on primary tumor biopsies and matched peripheral blood mononuclear cells (PBMCs), restricting to variants within the panel target regions (Figure 1e). cfDNA plasma was then subjected to deep targeted sequencing using the same hybrid-capture panel, and only reads overlapping catalog positions were retained for analysis. The validation cohort comprised 37 healthy controls and 126 stage I–IV CRC patients with pre-operative (pre-OP) plasma (median depth 8,367, IQR 9,225). The test cohort included 24 healthy controls, 209 stage I–III CRC patients, 29 minimally invasive pT1N0 cancers, and 17 adenomas (median depth 10,353, IQR 9,191 across the combined test and MRD cohorts). An additional 26 stage III patients provided post-operative (post-OP) plasma for MRD assessment (8/26 with confirmed relapse). All samples were collected, prepared, and processed as described by Frydendahl et al. [10].

Cancer detection was performed using a likelihood ratio test comparing a background-only model against a mixture model that allows a fraction *f* of tumor-derived ctDNA (Figure 1f). The test was applied across all read positions overlapping the patient’s mutation catalog. Samples with *f >* 0 were classified as ctDNA-positive and samples with *f* = 0 as ctDNA-negative.

### 2.2 cfDNA features

Two training datasets were produced: the SNV dataset contained 937,550 read-level mismatches and the indel dataset contained 128,483 read-level indels. For the SNV dataset, matches were randomly downsampled to achieve a 1:1 match-to-mismatch ratio. For the indel dataset, matches were downsampled to a 1:2 match-to-indel ratio for better balance across the three output classes (matches/SNVs, insertions, deletions).

Fragment size was associated with the estimated error rates for both SNVs and indels (Figure 2a). Error rates reached a minimum at fragment lengths close to the mononucleosomal peak (approximately 168 bp) and increased for shorter and longer fragments. Around the dinucleosomal peak, the error-rate curves did not show a reduction relative to surrounding fragment sizes, although the variance was lower in this region, consistent with higher read support around the dinucleosomal peak. Read counts showed a weak, approximately 10 bp periodicity between 100 and 160 bp, consistent with the helical pitch of nucleosomal DNA [28], and a corresponding modulation was observed in the error-rate curves over the same interval. The shortest fragments (approximately 50 to 70 bp) also exhibited a local reduction in estimated error rates relative to the immediately surrounding fragment-size range, for both variant classes, possibly reflecting protection by bound transcription factors.

**Figure 2.**
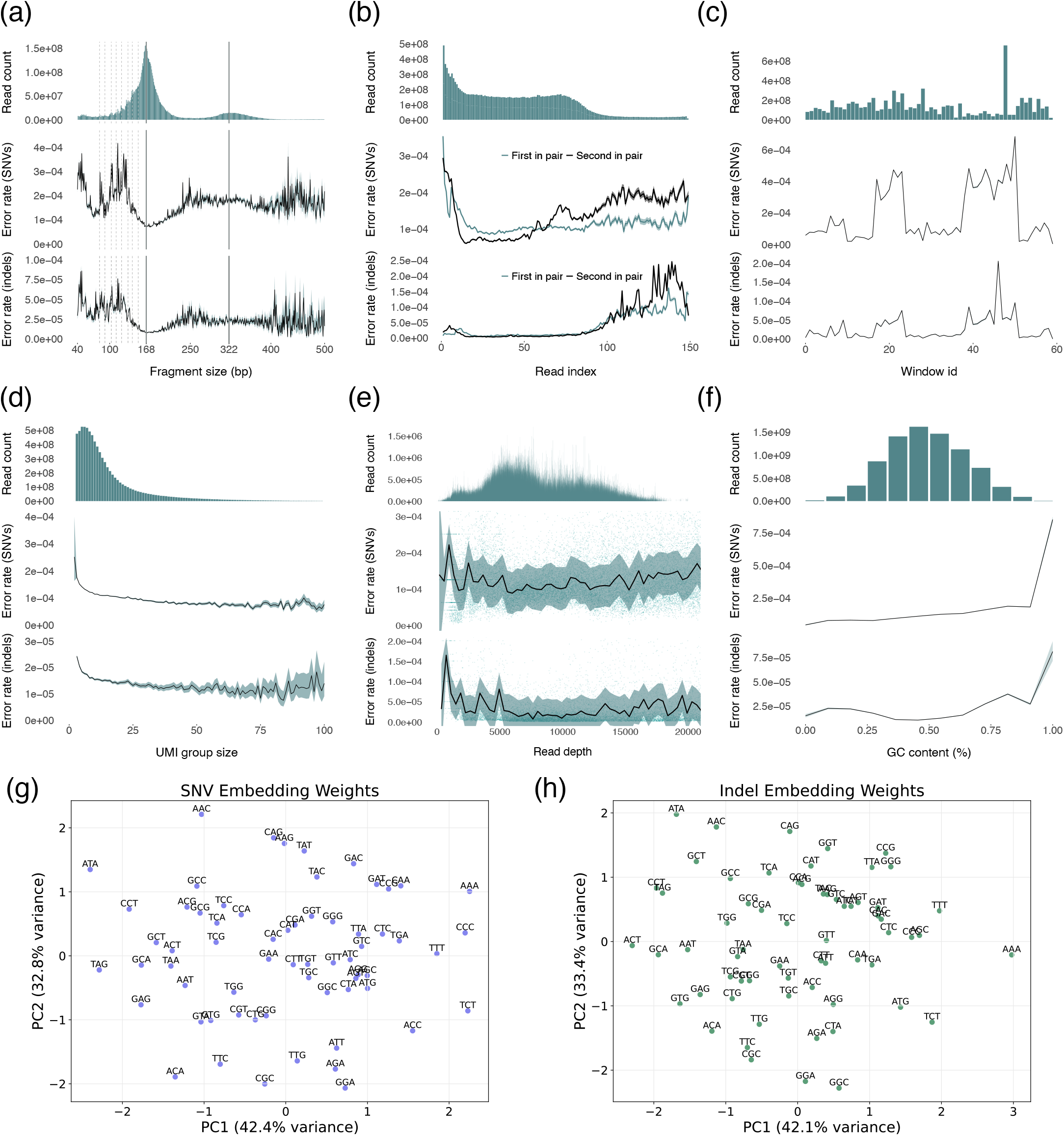
Systematic variation in cfDNA error rates across read and sequence features (a–f) Error rate as a function of read and sequence features. In each subpanel, the top plot shows the read count distribution; the middle and bottom plots show SNV and indel error rates, respectively. **(a)** Error rate vs fragment length. Error rates are lowest near the mononucleosomal peak (~168 bp), consistent with nucleosome-protected DNA being better preserved, and increase for shorter and longer fragments. Solid lines indicate the mononucleosomal (168 bp) and dinucleosomal (322 bp) peaks; dashed lines mark the ~10 bp periodicity reflecting the helical pitch of nucleosomal DNA. **(b)** Error rate vs read index, stratified by first and second read in pair. **(c)** Error rate vs window ID. **(d)** Error rate vs UMI group size. **(e)** Error rate vs read depth. Points represent individual genomic positions; lines show smoothed trends. **(f)** Error rate vs GC content. **(g–h)** PCA of learned trinucleotide embedding weights. Each point represents one of the 64 trinucleotide contexts, where proximity reflects learned similarity in error propensity. **(g)** SNV model. PC1 and PC2 explain 42.4% and 32.8% of variance, respectively. **(h)** Indel model. PC1 and PC2 explain 42.1% and 33.4% of variance, respectively.

Error rates as a function of sequencing cycle (Figure 2b) differed between SNVs and indels. SNV error rates were elevated during the first ~10 cycles, decreased to a broad minimum of approximately 1 × 10^−4^ across the read interior, and increased toward the 3^*′*^ end. In contrast, indel error rates remained close to zero across most cycles and then increased sharply after approximately cycle 100, with a spike near the read terminus, consistent with end-of-read base-quality decay and increased alignment ambiguity that affect indel calling more strongly than substitution calling. Error profiles also differed between the first and second read in pair across the full read length, with larger discrepancies for SNVs than for indels, consistent with a technical contribution from differences in sequencing chemistry and read orientation [30].

Across genomic windows (Figure 2c), read count varied substantially, with one window exceeding 6×10^8^ reads, which is suggestive of a highly repetitive locus or an amplicon-level imbalance. In some windows, SNV error rates were close to 1 × 10^−4^, but increased to approximately 7 × 10^−4^ in others. indel error rates showed a similar co-varying pattern at lower absolute rates. Together, these profiles indicate that a subset of windows contributes disproportionately to both depth and error, motivating inclusion of a window-level feature.

Error rates declined steeply with increasing UMI group size from three to approximately 10 reads and then plateaued (Figure 2d). SNV error rates decreased from roughly 3× 10^−4^ to about 1 × 10^−4^, whereas indel error rates stabilized near 1 × 10^−5^. The slight increase for very large groups coincided with wide uncertainty, suggesting sparse support rather than a systematic trend.

Read depth showed limited association with error rate across the well-sampled range (Figure 2e), consistent with read depth in hybrid-capture data reflecting variation in capture efficiency rather than an independent error determinant. From a few hundred to approximately 15,000 reads, SNV error rates remained near 1× 10^−4^ and indel error rates near 1 ×10^−5^.

Local GC content had only a modest effect on error rate through most of the range (Figure 2f). Read depth peaked around 40–60 % GC and fell toward both extremes. SNV errors stayed near 1 × 10^−4^ through much of the distribution, rose gradually with increasing GC and surged to 9× 10^−4^ in the highest bin, where counts were sparse. Indel errors followed the same trend at roughly one-tenth the magnitude. The surge in error rates at high GC coincided with sparse coverage, suggesting that GC-related difficulties in amplification and alignment both reduce coverage and drive error rates upward, making GC content a strong predictor of error specifically at the extreme high end of the distribution.

To visualize the learned trinucleotide representations, principal component analysis (PCA) was applied to the embedding weights of the SNV model (Figure 2g). The first two principal components explained 42.4% and 32.8% of the variance in the embedding weights, respectively. Each point represents one of the 64 trinucleotide contexts, where nearby points contribute similarly to predicted error propensity after accounting for other features. The PCA reveals gradual structure without discrete clusters.

The indel model embedding produced a distinct layout under PCA (Figure 2h), with PC1 and PC2 capturing 42.1% and 33.4% of the variance, respectively. The arrangement differed from the SNV embedding, indicating that the trinucleotide contexts most predictive of indel errors are not the same as those for SNVs.

### 2.3 Neural network model of SNV and indel error rates

Two separate neural networks were trained: one to predict SNV error rates and one to predict indel error rates (Figure 3a) (Methods, Section 5.3.1). The neural network models took as input numeric features, categorical features, and a single embedded feature (Methods, Section 5.3.3). The full model contained all features depicted in Figure 3a. Both network architectures were optimized separately using Weights & Biases (W&B) Sweeps [2] for hyperparameter optimization with the full feature set (Appendix, Figure A1 for SNVs, Figure A2 for indels, and Table A1).

**Figure 3.**
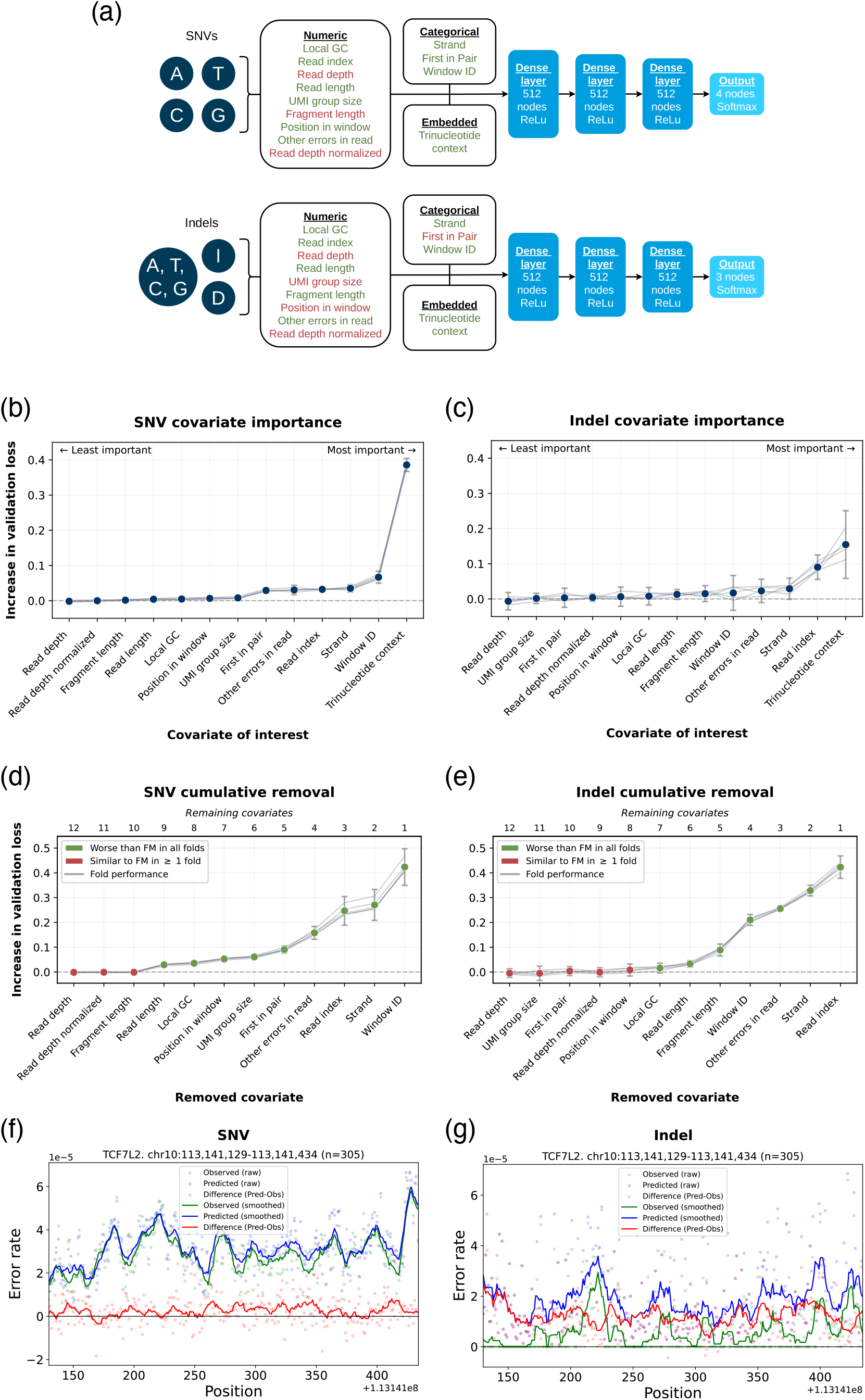
Neural network models for SNV and INDEL error rates. **(a)** Model architectures. features are divided into three categories: numeric, categorical, and embedded. Features in transparent were omitted in the final model. **(b–c)** feature importance. Quantified as the increase in validation loss when a feature is removed from the full model (LOCO). **(d–e)** Cumulative feature removal. Models showing the change in validation loss as features are removed sequentially in LOCO order. Green indicates improved performance relative to the full model, and red indicates worse performance. **(f–g)** Predictions vs observations at the TCF7L2 locus. In a held-out validation region (chr10:113,141,129 to 113,141,434; *n* = 305 positions), the prediction performanace was evaluated for both SNVs **(f)** and indels **(g)**. Points represent perposition error-rate estimates aggregated across reads and corrected for match downsampling; solid lines show Savitzky-Golay smoothed trends. Green denotes observed error rates, blue denotes model predictions evaluated at the same positions, and red denotes residuals (prediction minus observation).

To quantify relative feature importance, remove uninformative features, and limit model complexity, we iteratively evaluated feature contributions in a leave-one-covariate-out (leave one covariate out (LOCO)) approach, following Lei et al. [14]. A baseline model containing all features was first evaluated, then restricted models were fit, each omitting one feature in turn. The change in performance relative to the baseline was used to measure and rank feature importance (Figure 3b–c). For SNVs, the dominant signal came from trinucleotide context, which yielded by far the largest increase in validation loss when omitted (Figure 3b). Secondary effects were observed for window ID and read index, with smaller but nontrivial contributions from strand, other errors in the read, and whether the read was first or second in pair. Read depth, normalized depth, fragment length, read length, local GC content, and position in window had negligible impact, presumably because they account for only a small fraction of the predictive signal once trinucleotide context and read-level features are already included. For indels, the most informative feature was also trinucleotide context, followed by modest effects of read index, strand, and other errors in read (Figure 3c). Notably, indel error rates are very low (typically below 5 × 10^−5^) across most read positions and spike sharply only at specific late-cycle indexes, making read index a particularly informative covariate for indels despite its moderate overall importance. Features such as read depth, UMI group size, and first in pair had little to no effect. These rankings reflect conditional drops in performance and may be dampened or inflated by collinearity among features; they should therefore be interpreted as feature-to-loss correlations under the full model rather than as independent, additive contributions [14, 8].

Next, an informative subset of features was selected by backward stepwise elimination. Starting from the full feature set, features were removed one at a time in LOCO order while tracking the change in validation loss (Figure 3d–e). The final subset was the smallest model whose performance did not degrade relative to the full model. For SNVs, removing normalized read depth, read depth, and fragment length had negligible or slightly positive effects, so these were discarded. Performance began to degrade after removing read length and local GC content, and dropped further upon eliminating position in window, UMI group size, first in pair, other errors in the read, read index, strand and especially window ID (Figure 3d). For indels, read depth, UMI group size, first in pair, normalized read depth, and position in window could be removed with no increase in validation loss (Figure 3e). Validation loss began to rise after removing local GC, read length, fragment length, window ID, other errors in the read, and strand, with the largest penalty observed when dropping the read index. Accordingly, for both SNVs and indels, the final model was chosen as the smallest feature subset whose validation loss was no worse than that of the full model.

To assess calibration at nucleotide resolution, we aggregated read level observations by genomic coordinate and plotted position-wise error rates after correcting for deliberate down-sampling of matches (Figure 3f and 3g).

In the SNV window (Figure 3f), the predicted and observed smoothed error rate curves closely track each other across the interval, indicating good calibration of the broad position-wise structure. Residuals (prediction minus observation) remain small and centered near zero, with modest, locally correlated deviations around sharper peaks and troughs, suggesting residual position-specific effects that are not fully explained by the current features.

In the indel window (Figure 3g), observed error rates are lower overall and more sporadic. The model reproduces several local increases but tends to predict higher baseline rates than observed, yielding predominantly positive residuals across much of the region. The largest discrepancies occur at a subset of positions with pronounced spikes, consistent with sparse events and remaining unmodeled determinants of indel error, such as low-complexity se-quence, local alignment ambiguity, or context around indel-prone motifs.

### 2.4 Statistical model to separate signal from noise

To detect ctDNA, reads overlapping each position in the patient-specific mutation catalog are interrogated to determine whether the observed mismatches are better explained by background sequencing error alone or by a mixture of tumor-derived signal and error. Following the framework of Christensen et al. [6], each candidate mutation site — a genomic position from the patient’s catalog — was evaluated with a likelihood-ratio test. Under the null hypothesis, the site was assumed absent and observed mismatches were attributed to sequencing and alignment errors (*H*_0_), whereas the alternative assumed the site was present heterozygously and reads followed a mixture of reference and mutant alleles driven by the tumor fraction (*H*_1_). A trained neural network mapped read-level features to per-read error rates, which parameterized the observation model and entered the likelihoods under both *H*_0_ and *H*_1_. Mutational ctDNA evidence was summarized by the test statistic *Q* = −2 log(*L*_0_*/L*_1_) (Figure 4a).

**Figure 4.**
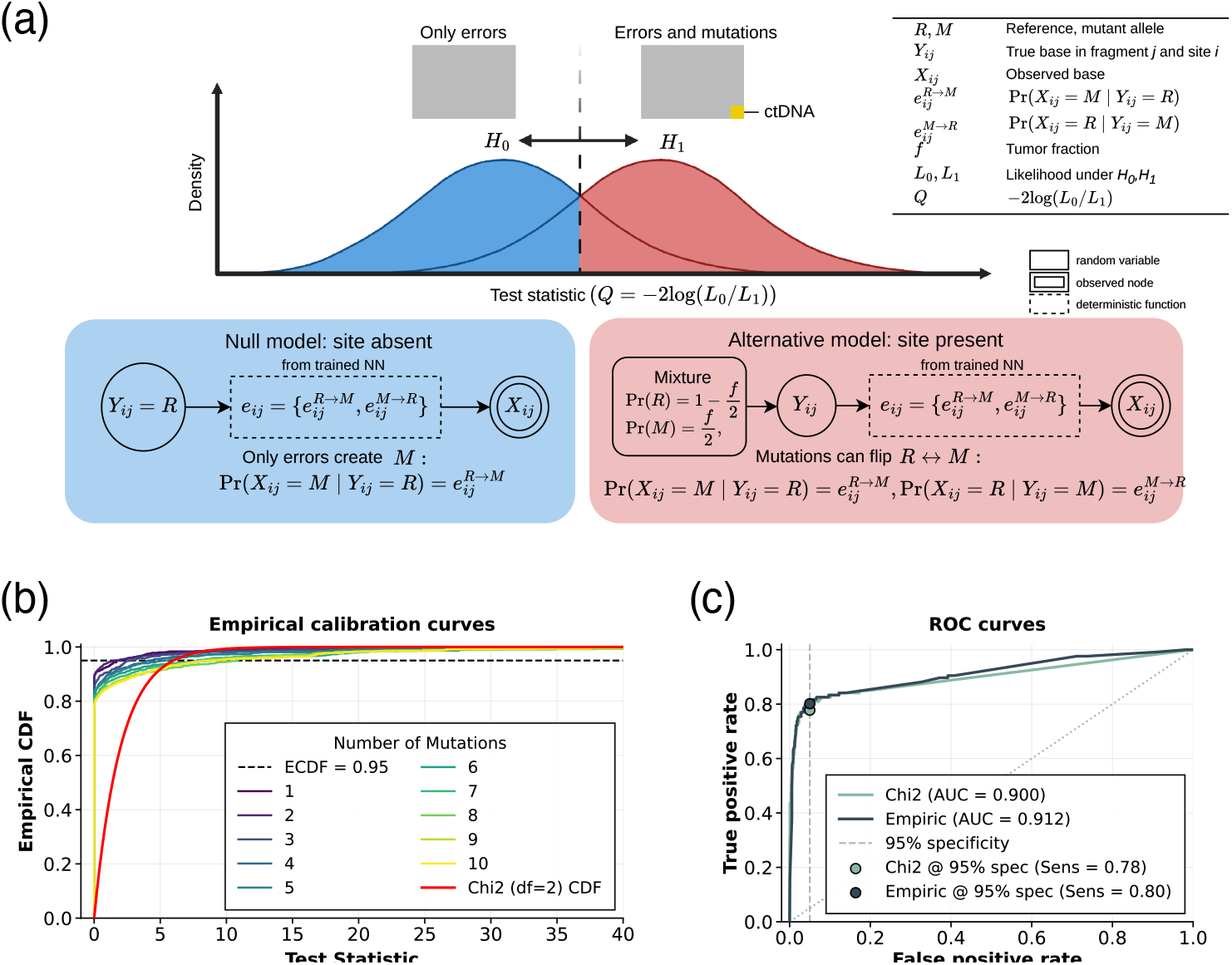
Statistical model for ctDNA detection and calibration. **(a)** Probabilistic graphical model used at calling time. For each candidate site *i*, the latent variable *Z*_*i*_ indicates whether the mutation is present. For each read *j* at that site, *Y*_*ij*_ *∈* {*R, M*} is the latent true allele and *X*_*ij*_ is the observed allele. A trained neural network maps read-level features *D*_*ij*_ to per-read error rates, which parameterize the likelihood of the observed allele under both the null (*Z*_*i*_ = 0, errors only) and the alternative (*Z*_*i*_ = 1, mixture of tumor and reference alleles at fraction *f*). Global parameters are the site-presence probability *r* and the tumor fraction *f*. Multi-class network outputs are collapsed to a binary likelihood for SNV and indel candidates. (b) Empirical calibration curves based on 12,000 pseudo-healthy controls with known ctDNA-negative status, generated by randomly pairing 46 healthy controls with randomly shuffled mutation catalogs. Calibration curves are stratified by mutation catalog size. **(c)** ROC curves comparing ctDNA detection performance under empirical calibration (thresholds estimated from the observed null distribution of *Q*) versus the *χ*^2^ approximation, evaluated in the validation cohort of CRC patients and healthy controls.

To classify a sample as ctDNA-positive or -negative, *Q* must be compared against a threshold corresponding to a desired specificity. Because the null distribution of *Q* may deviate from its asymptotic approximation in practice, the choice of calibration strategy can affect the operating point. Two calibration strategies were therefore evaluated: a *χ*^2^ approximation as done by [6] and an empirical calibration based on the observed null distribution of *Q* derived from healthy-control samples (Figure 4b). The empirical approach is motivated by the observation that *Q* does not follow a *χ*^2^ distribution in practice, making data-derived thresholds more appropriate. Empirical calibration yielded improved discrimination relative to the *χ*^2^ approximation, with an AUC of 0.912 compared with 0.900 — a ~10% relative reduction in classification error (Figure 4c). At the 95% specificity operating point, empirical calibration also increased sensitivity (0.80 vs 0.78). The corresponding thresholds were p-value 0.0395 for the empirically calibrated test statistic and p-value 0.307 for the *χ*^2^-based test statistic (Figure 4b–c). The large difference in thresholds reflects the fact that *Q* is overdispersed relative to the *χ*^2^ distribution — the *χ*^2^ approximation requires a much more lenient p-value cutoff to achieve the same nominal specificity.

### 2.5 Contribution of SNVs and indels to ctDNA detection

To quantify the relative contribution of each variant class, detection performance was evaluated using three call sets: SNVs only, indel only, and the combined set (SNVs + indels) in the validation cohort.

To ensure a fair comparison between call sets, the analysis was restricted to patients for whom both SNVs and indels were present in their mutation catalog, so that all three call sets could be evaluated in the same individuals. This applied to the majority of the validation cohort: 66 of 126 CRC patients (52%) across all stages, together with 37 healthy controls (*n* = 66 pre-OP CRC: stage I: 16, stage II: 31, stage III: 13, stage IV: 6; *n* = 37 controls) (Figure 1a). Mutation catalogs from the 66 cancer patients were randomly permuted and reassigned to controls to generate 66 catalog-matched healthy individuals (Methods, Section 5).

To assess the per-mutation evidence provided by each variant class, PyDREAMS was run twice per patient — once using only SNV loci and once using only indel loci — and the resulting variant-type-specific score −log_10_(*p*) was divided by the number of mutations of the corresponding type in the patient’s tumor-informed panel (Methods, Section 5.4). Indels contributed a higher score per mutation than SNVs (median ratio indel/SNV ~1.5; Figure 5a), indicating that, on average, a single indel provides more evidence for ctDNA presence than a single SNV under the DREAMS model.

**Figure 5.**
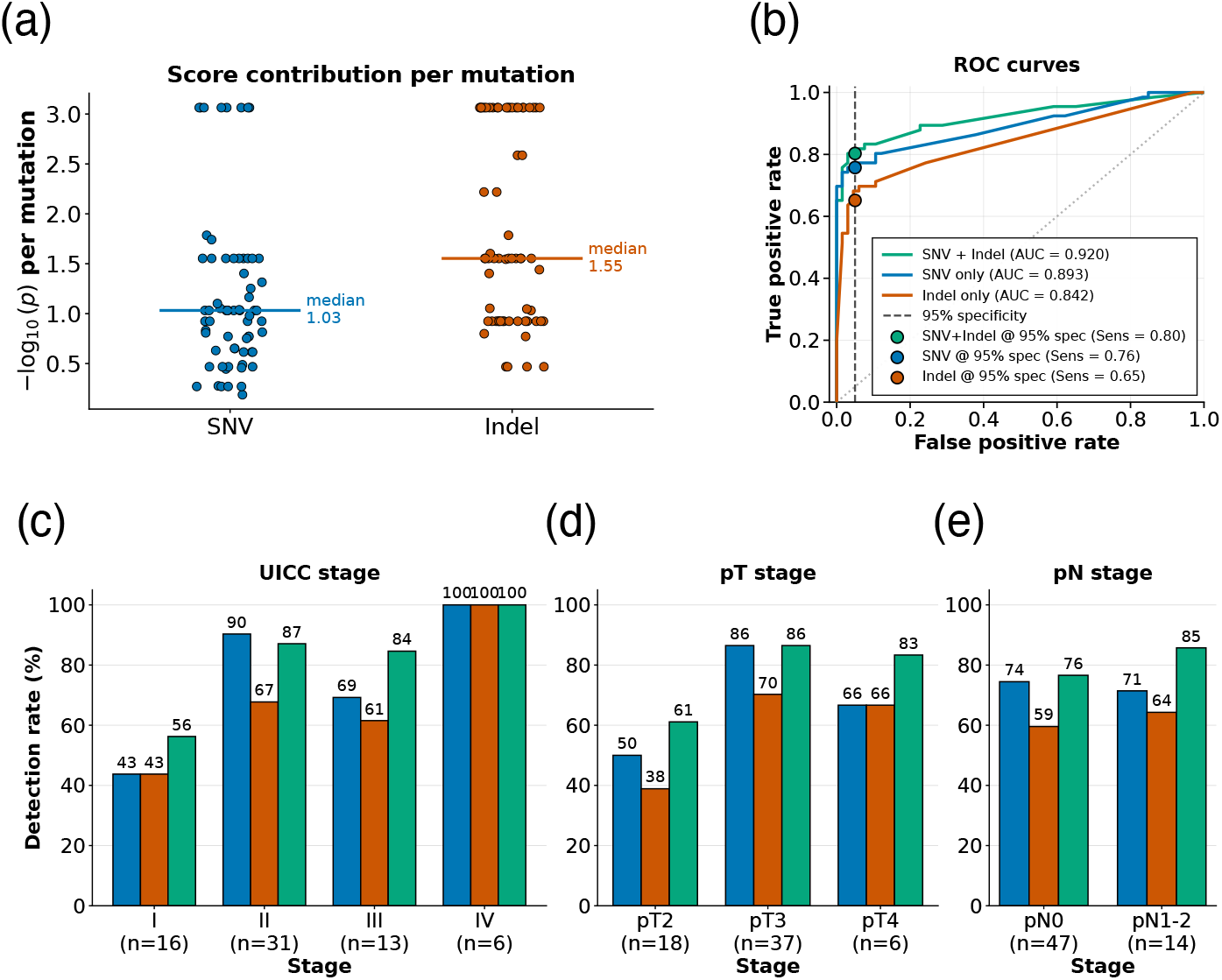
ctDNA detection performance in the validation cohort. **(a)** Per-mutation score contribution for SNVs and indels across the CRC cohort, shown as a paired scatter plot with each patient represented by an SNV and an indel point. The per-mutation contribution is defined as − log_10_(*p*)*/n*, where *p* is the variant-type-specific p-value obtained by running PyDREAMS with SNV-only or indel-only loci, and *n* is the number of mutations of the corresponding type in the patient’s tumor-informed panel. Horizontal bars mark the median permutation score for each variant type, highlighting the difference in means. Only patients carrying at least one SNV and at least one indel are included. **(b)** ROC curves. Contribution of SNVs and indels to detection performance, evaluated using SNVs only, indels only, and the combined call set (SNVs + indels). The subset shown includes individuals for whom both SNVs and indels were present in the mutation catalog (66 cancer patients and 37 healthy controls; mutation catalogs for the 66 cancer patients were randomly permuted and reassigned to controls to generate 66 catalog-matched healthy individuals). **(c)** UICC stage. **(d)** pT stage. **(e)** pN stage.

The combined call set (SNVs + indels) achieved the best discrimination (area under the curve (AUC) 0.920), outperforming SNV-only (AUC 0.893) and indel-only (AUC 0.842) (Figure 5b). At 95% specificity, the combined call set also achieved the highest sensitivity (0.80), followed by SNV-only (0.76) and indel-only (0.65) (Figure 5b).

At 95% specificity, detection rates generally increased with more advanced disease, although not strictly monotonically across stages (Figure 5c). For UICC stage I, detection was 43% for SNV-only, 43% for indel-only, and 56% for the combined call set. For stage II, detection was 90%, 67%, and 87%, respectively. For stage III, detection was 69%, 61%, and 84%, and for stage IV all three call sets reached 100% (noting the small sample size for stage IV, *n* = 6).

Stratification by pathological t (pT) and nodal (pathological n (pN)) stage showed that adding indels to SNVs consistently improved or matched detection across subgroups (Figure 5d–e). The gain from combining variant classes was most apparent in pT2 (combined: 61% vs SNV-only: 50%) and pN0 (combined: 85% vs SNV-only: 76%), whereas differences were negligible in pT3 and pN1-2.

### 2.6 ctDNA detection performance in the validation cohort

Next, PyDREAMS was benchmarked against UMIseq, the method previously applied to this cohort by Frydendahl et al. [10], making it the natural reference for comparison. UMIseq is a Shearwater-based variant caller that estimates position-specific error rates from a panel of normals, representing an established approach against which the neural network-based error model of PyDREAMS is evaluated. The benchmark was first performed in the validation cohort (Figure 1a), which was also used to calibrate both methods by selecting a method-specific decision threshold from its receiver operator characteristics (ROC) curve. In the validation cohort, 100 of 121 patients (82.64%) had detectable ctDNA signal, defined as at least one read supporting a catalog variant observed in the cfDNA data; all reported results include the full cohort, including the 21 patients without detection. Overall discrimination was similar between methods, with PyDREAMS achieving a slightly higher AUC than UMIseq (0.917 vs 0.909; Figure 6a). At the predefined operating point of 95% specificity, PyDREAMS also showed marginally higher sensitivity than UMIseq (0.82 vs 0.80; Figure 6a).

**Figure 6.**
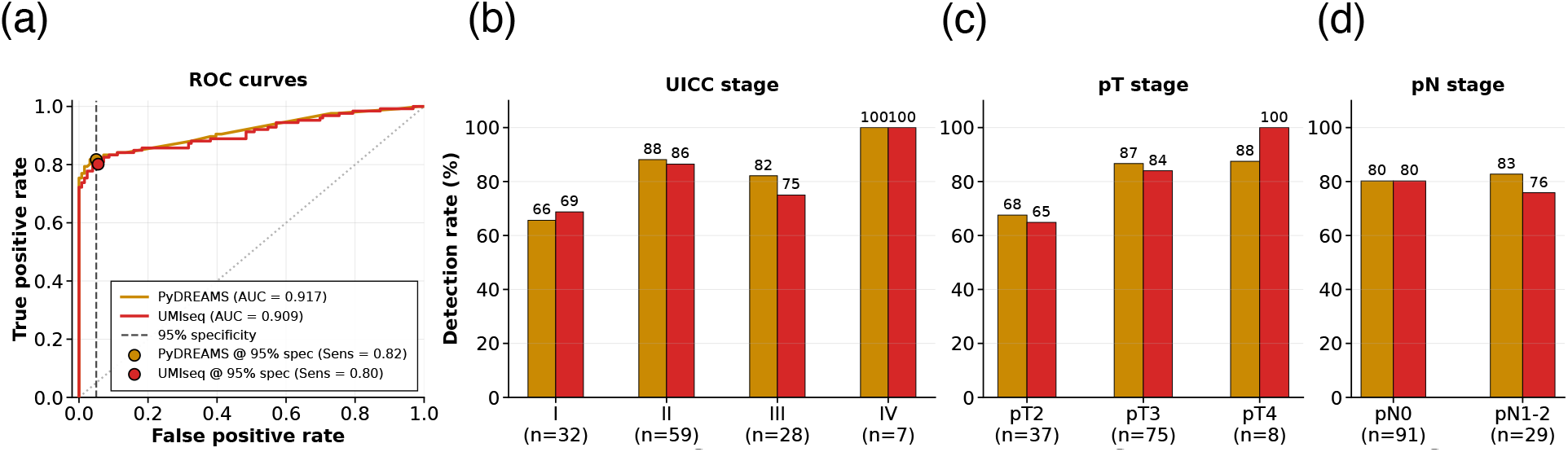
ctDNA detection in the validation cohort. **(a)** ROC curves. **(b)** UICC stage. **(c)** pT stage. **(d)** pN stage.

Detection rates stratified by clinicopathological variables were broadly comparable between methods (Figure 6b to d). By UICC stage, detection increased with advancing stage for both methods, ranging from 66% (PyDREAMS) and 69% (UMIseq) in stage I to 100% for both in stage IV (Figure 6b). Stage II showed high detection for both methods (88% vs 86%), while PyDREAMS was higher in stage III (82% vs 75%; Figure 6b). By pT stage, detection was 68% vs 65% in pT2 and 87% vs 84% in pT3, while UMIseq was higher in pT4 (100% vs 88%; Figure 6c). By nodal stage, detection was identical in pN0 (80% for both) and higher for PyDREAMS in pN1-2 (83% vs 76%; Figure 6d).

### 2.7 ctDNA detection performance in the test cohort

ctDNA detection performance was compared between PyDREAMS and UMIseq in the Roche test cohort (Figure 7), which comprises three distinct subgroups: a main CRC tumor cohort (stage I–III, *n* = 209), an early-stage and adenoma subset (pT1pN0 and adenomas, *n* = 46), and a post-OP MRD subset (*n* = 26). Overall discrimination was assessed in the main tumor cohort only, as the early-stage and MRD subsets represent separate clinical questions analyzed below. PyDREAMS achieved a slightly higher AUC than UMIseq in this cohort (0.917 vs 0.909; Figure 7a). At the predefined operating point of 95% specificity, the sensitivity was 0.82 for PyDREAMS and 0.80 for UMIseq (Figure 7a). In this main tumor cohort, detectable ctDNA signal was observed in 196 of 209 samples (93.8%).

**Figure 7.**
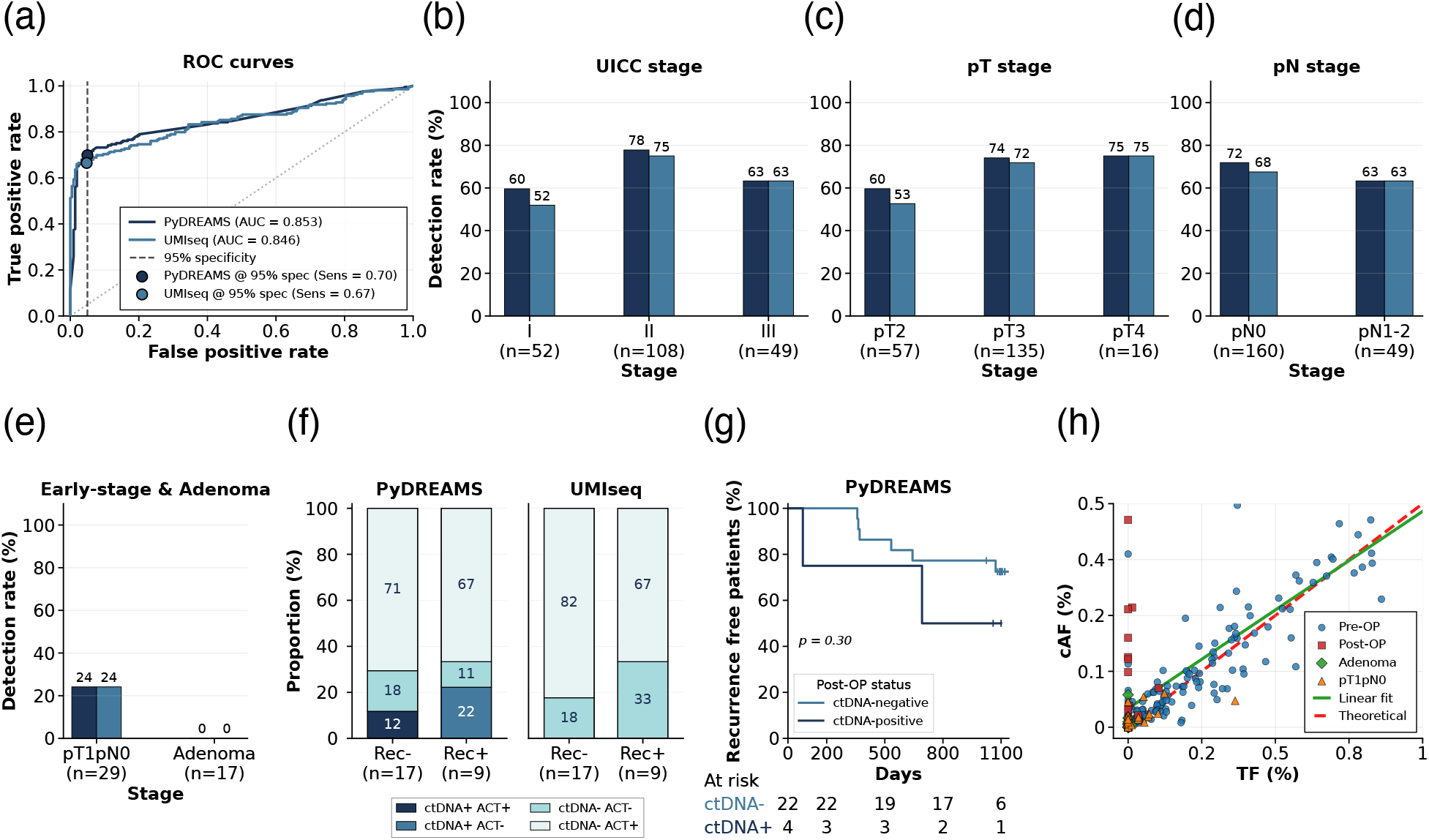
ctDNA detection performance across clinical subgroups in the test cohort. **(a)** ROC curves. **(b)** UICC stage. **(c)** pT stage. **(d)** pN stage. **(e)** Early-stage & Adenoma. **(f)** MRD detection in post-OP samples. **(g)** Kaplan Meier analysis for PyDREAMS. post-OP samples were used. **(h)** Tumor fraction vs circulating tumor allele frequency.

Stratified analyses showed comparable detection rates across clinicopathological sub-groups within the main tumor cohort (Figure 7b–d). By UICC stage, PyDREAMS showed higher detection in stage I (62% vs 54%) and stage II (81% vs 77%), while performance was identical in stage III (67% for both) (Figure 7b). By pT stage, PyDREAMS again showed higher detection in pT2 (61% vs 54%) and pT3 (78% vs 74%), with identical detection in pT4 (81% for both; Figure 7c). By nodal stage, PyDREAMS showed higher detection in pN0 (75% vs 69%), whereas detection was identical in pN1-2 (67% for both; Figure 7d).

Performance was further evaluated in early-stage disease and adenomas (Figure 7e). In pT1pN0 patients, both methods detected ctDNA in 34% of cases (*n* = 29), despite tumor mutations being present in the cfDNA data in 82.8% of samples. No adenoma samples were classified as ctDNA-positive by either method (*n* = 17).

Finally, PyDREAMS was evaluated in the MRD setting, where post-OP ctDNA detection may identify patients at risk of recurrence before clinical relapse becomes apparent. This analysis was restricted to the subset of stage III patients with available post-OP plasma and recurrence follow-up (*n* = 26, of whom 9 later relapsed; Figure 7f–g). At least one catalog tumor mutation was detectable in the cfDNA data in 5 of 26 post-OP samples (19.2%). PyDREAMS classified 5/26 patients as ctDNA-positive post-OP, including 3/9 patients who later recurred and 2/17 patients who remained recurrence-free, corresponding to an enrichment for recurrence of (3*/*5)*/*(9*/*26) = 1.73 among ctDNA-positive patients relative to the cohort baseline recurrence rate. In contrast, UMIseq did not classify any post-OP samples as ctDNA-positive in this subset (Figure 7f). Kaplan Meier analysis for PyDREAMS showed lower recurrence-free survival for post-OP ctDNA-positive patients, approaching near-significance (log-rank *p* = 0.06; Figure 7g).

Across samples, PyDREAMS tumor fraction estimates were positively associated with the observed circulating allele frequency (cAF) — the fraction of circulating alleles carrying a tumor mutation — and the empirical linear fit closely followed the theoretical relationship at higher tumor fractions (Figure 7h). Early-stage (pT1pN0) and adenoma samples clustered at low tumor fractions and low cAF, consistent with minimal tumor shedding into the circulation and explaining the limited detection rates in these subgroups.

## 3 Discussion

PyDREAMS extends the DREAMS read level error modeling framework from SNVs to indels, enabling unified tumor informed ctDNA detection in settings where patient specific catalogs contain a mix of variant classes, or where indels contribute substantially to tumor signal. We exemplify this for CRC, where clinically informative alterations include both SNVs and indels, and where the low ctDNA fractions encountered in early stage disease and MRD place stringent demands on background modeling. Using UMI collapsed consensus reads, PyDREAMS aims to explain residual technical noise that persists after consensus building, and to translate read level error probabilities into calibrated evidence for or against tumor derived signal. It is also provided as an easy-to-use, installable Python package, lowering the barrier to adoption in other targeted cfDNA sequencing studies.

Fragment size emerged as an error-informative feature for both SNVs and indels, revealing a link between cfDNA protection states and residual sequencing background. The same fragment-length structure that governs SNV error rates (as shown previously [6]) also holds for indels. Interestingly, for both SNVs and indels, the lowest error rates occurred precisely at fragment sizes corresponding to mononucleosomal lengths, directly linking nucleosome protection to reduced residual error and suggesting that nucleosome-wrapped fragments are not only more stable in circulation but also better protected from lesions, leading to fewer apparent sequencing errors either during circulation or sample handling. Fragments outside the nucleosome-protected size range were less abundant and more error-prone, generally consistent with increased nuclease-mediated degradation during circulation [13, 26, 27].

Interestingly, very short fragments (approximately 50 to 70 bp) also showed reduced error rates for both SNVs and indels. This suggests that this length range may represent a distinct, more protected fragment class, plausibly enriched for genomic regions protected by bound proteins such as transcription factors (TFs), which could reduce nuclease accessibility and thereby limit the accumulation of damage-associated artifacts. This interpretation is plausible given prior reports that cfDNA fragmentation patterns encode TF footprints [28, 32, 24]. More generally, extending the error model with TF-aware features, such as bindingsite overlap or footprint strength, may improve error-rate calibration, particularly for short fragments.

The sequencing-cycle profiles differed between SNVs and indels, suggesting distinct dominant error sources for the two variant classes. Elevated SNV errors in the first cycles and toward the 3^*′*^ end are consistent with damage at fragment termini — such as oxidative lesions and end-repair artefacts introduced during library preparation — with a possible secondary contribution from cycle-dependent base-calling quality decay due to clusters going out of phase across the flowcell [29]. In contrast, the late-cycle rise in indel errors may reflect a combination of increased base-calling uncertainty at low-quality read ends and increased alignment ambiguity, where gapped alignments can be misplaced, particularly in repetitive or low-complexity sequences [18, 7]. Disentangling read-level from alignment-level contributions could be approached by inspecting partially overlapping read pairs, where the same molecule is sequenced from both ends. The systematic differences between the first and second read in pair further support a technical contribution related to sequencing chemistry and read orientation [30]. Together, these patterns motivate modeling error rates as a function of cycle position and read in pair, and treating SNVs and indels separately to avoid miscalibration in low-frequency variant calling.

A plausible explanation for the observed window-specific spikes in read count and, separately, elevated error rates is local genomic context. Highly repetitive or duplicated loci can attract ambiguously mapped reads, inflating apparent depth, while also increasing apparent SNV and indel error rates through misalignment, context-specific artefacts, and the cumulative effects of errors introduced over many repeated PCR cycles. Window-specific error spikes could further arise where target sequence composition concentrates Illumina artefacts — for example, homopolymer runs that drive phasing/pre-phasing errors and indel calling, or stretches that produce low-chastity clusters failing signal-purity filters. These mechanisms could also help explain the elevated cycle-late indel error rates discussed above. Importantly, these effects need not co-occur within the same windows: some loci may primarily drive depth through multi-mapping, whereas others may primarily drive errors through alignment ambiguity, sequence composition, or indel-prone contexts [31, 22].

As expected, UMI-based consensus calling substantially reduces error rates — approximately 3-fold for SNVs and up to 10-fold for indels — with diminishing returns beyond roughly 10 reads per group.

Local GC content had little impact on error rates across the well covered range, but errors increased sharply in the most GC rich bin where coverage was sparse. This suggests GC becomes informative mainly at the extreme, likely reflecting a combination of GC related coverage dropouts and increased technical difficulty in amplification and mapping [4]. Hence, it may be useful to include local GC content as a feature in the error model, particularly to improve calibration at GC-extreme positions.

The neural networks for both SNVs and indels were first optimized via a hyperparameter sweep over network width, depth, learning rate, batch size, and dropout rate, identifying a configuration — three hidden layers of 512 units each with dropout — that reduced validation loss relative to the original DREAMS architecture [6]. In both cases, the hyperparameter search identified a three hidden-layer architecture, consistent with the initial design, but with higher capacity per layer. Although deeper and wider configurations could also achieve strong performance, the best configuration among the evaluated runs used three layers of 512 units each ([512, 512, 512]). A plausible explanation is a bias-variance and optimization trade-off: increasing depth and width raises expressive power but can reduce training stability and increase overfitting given noisy targets and finite effective sample size. Under these conditions, moderate capacity can generalize better than more complex alternatives within a finite hyperparameter search. On larger training sets, a larger network may become more attractive, and users with access to more healthy-control samples should consider re-running the hyperparameter sweep accordingly.

LOCO shows that trinucleotide context dominates for both SNVs and indels, while depth-related features (read depth and normalized depth) add little conditional information. The residual contributions of read index, strand, and “other errors in read” are consistent with cycle-dependent, strand-asymmetric artifacts and within-read error correlation. The prominence of window ID, especially for SNVs, also suggests that region-specific heterogeneity is an informative feature. Because LOCO is conditional, importances should be read as pragmatic under collinearity rather than causal [14, 8].

Backward elimination supports a compact feature set — depth-derived features can be removed without loss, while SNVs remain sensitive to window-level effects and indels to read index. Nucleotide-resolution calibration confirms this: SNV residuals are small and localized, suggesting missing fine-scale interactions, whereas indel residuals show an elevated baseline and occasional large spikes, consistent with a sparse, heterogeneous process shaped by unmodeled sequence and alignment determinants.

Empirical calibration of the likelihood-ratio statistic is a key methodological contribution of PyDREAMS. Discrimination performance is driven not only by the likelihood-ratio formulation, but also by how its null distribution is calibrated in the low-count, low allele-fraction regime relevant for ctDNA. The modest but consistent gain from empirical calibration suggests that the asymptotic *χ*^2^ approximation is imperfect under the practical conditions of the test — as is also directly visible in the test-statistic-to-p-value calibration curves (Figure 4b) — for example due to boundary constraints (presence versus absence), model misspecification in the error parameterization, or dependence and heterogeneity across reads. In this setting, learning the null distribution directly from data yields thresholds that better match the realized Type I error, which in turn improves sensitivity at a fixed specificity.

The large difference in p-value cutoffs reflects meaningfully different tail behavior of *Q*, highlighting a failure mode of asymptotic theory for operating-point selection in ultra-low-frequency variant detection.

Jointly calling SNVs and indels improves ctDNA detection because it increases the number of informative loci, averages over class-specific noise processes, and incorporates indels, which have inherently lower background error rates and therefore contribute disproportionately informative signal per locus. Consistent with this, the per-mutation score contribution (Figure 5a) showed that, on a per-locus basis, a single indel provided ~1.5× more evidence for ctDNA presence than a single SNV, directly reflecting the lower background error rate at indel loci. The stronger performance of the combined call set, relative to either class alone, is therefore consistent with a simple signal-aggregation effect rather than a single class being uniformly superior. The weaker indel-only performance likely reflects the higher heterogeneity and sparsity of indel observations, which inflates variance at fixed specificity. The stage-stratified results support the expected dependence on tumor burden, but the non-monotonicity across stages and subgroups is plausibly driven by modest sample sizes within strata and biological heterogeneity in shedding, rather than a systematic failure of the classifier.

PyDREAMS still requires representative training data, but replaces locus-specific PON error estimation with a transferable, feature-based model. Performance differences are small and within the range attributable to cohort composition and threshold calibration. Note that 21 of 121 patients (17%) had no detectable ctDNA signal; retaining these samples means reported sensitivities reflect a clinically realistic mixture of detectable and non-detectable cases.

The test cohort results reinforce this picture and argue against the validation performance being purely cohort-specific. In the main tumor cohort, where the AUC comparison was computed, detectable ctDNA signal was present in 196 of 209 samples (93.8%), indicating that the observed discrimination largely reflects separation among predominantly signal-positive cases rather than robustness to pervasive non-detectability. At the same time, signal prevalence dropped markedly in lower-signal settings, with detectable ctDNA observed in 24 of 29 pT1pN0 samples (82.8%), 6 of 17 adenoma samples (35.3%), and 5 of 26 post-OP samples (19.2%). The comparable subgroup trends, together with the absence of ctDNA-positive calls in adenomas at the selected threshold, are consistent with controlled specificity even when measurable signal is uncommon, although the presence of detectable signal in a subset of adenomas highlights that background signal alone is not sufficient to trigger positivity under the applied decision rule. The post-OP analysis is suggestive but should be treated as exploratory: ctDNA-positive calls occurred in a minority of samples despite low signal prevalence overall, and their enrichment among later recurrences aligns with the expected direction of effect, yet the limited sample size and borderline log-rank result motivate evaluation in larger postoperative cohorts with prespecified thresholds, standardized sampling time points, and longer follow-up

PyDREAMS is released as an open-source Python package with documented workflows and reproducible dependency management, facilitating benchmarking, adaptation, and reuse in targeted cfDNA sequencing studies.

Finally, the positive association between estimated tumor fraction and observed cAF provides an internal consistency check on the quantitative output of PyDREAMS. Deviations at low tumor fractions are expected because sampling noise and residual model misspecification dominate when true signal approaches the detection limit, which is also the regime where differences between callers are most sensitive to calibration choices. More broadly, a large fraction of apparent mismatches in cfDNA sequencing likely originate from DNA lesions — such as oxidative damage and deamination — that are converted into fixed mutations during PCR-based library preparation, rendering them indistinguishable from true somatic variants. Recognizing this as a dominant source of background error, rather than random sequencing noise, has important implications for how error models should be designed and interpreted across the field.

## 4 Conclusion

PyDREAMS extends read-level cfDNA error modelling from SNVs to indels, providing a unified, tumor-informed background error model for mixed mutation catalogs in deep targeted plasma sequencing. By revealing that cfDNA fragment structure and sequencing-cycle effects shape residual background error rates for both variant classes, PyDREAMS connects biological cfDNA features to technical noise in a way that goes beyond descriptive fragmentomics. Distinct error profiles for SNVs and indels justify separate neural network models and show that the two variant classes cannot be treated under a single generic noise process. Empirical calibration of the likelihood-ratio statistic improves operating-point control in the ultra-low allele-fraction regime, where asymptotic approximations are unreliable. Jointly modelling SNVs and indels improves ctDNA detection relative to either class alone, and PyDREAMS achieves performance comparable to, or slightly better than, existing error-modelling approaches, with suggestive benefit in postoperative low-signal samples relevant to MRD assessment. In summary, PyDREAMS extends read-level cfDNA error modelling from SNVs to indels and shows that fragment-informed, empirically calibrated background modelling can improve the interpretation of ultra-low-frequency tumor signal in deep targeted plasma sequencing. Released as an open-source Python package, PyDREAMS facilitates benchmarking, adaptation, and reuse across targeted cfDNA sequencing studies.

## 5 Methods

### 5.1 Read-level error-rate prediction for SNVs and indels

Per-base sequencing error rates were estimated using the read-level error-modelling approach described in DREAMS [6]. The method was reimplemented in Python and extended to support indels in addition to SNVs. For each sequenced locus, per-read error probabilities were predicted using separate neural network models for SNVs and indels (Figures 1c and 3a).

Data and feature vectors were derived from UMI-collapsed consensus reads in BAM files generated by next-generation sequencing, as described in [6]. Models were trained on the training cohort and applied unchanged to the validation, test, and MRD cohorts. Read mapping, consensus generation, and preprocessing followed [10]. Class imbalance between matches and mismatches was handled as described in [6]. In addition to the features used in DREAMS, window ID, read depth, and normalized read depth were included; window ID is a unique identifier for each target genomic region defined in the input BED file.

Read depth normalization was performed on a per-window basis to account for variation in target capture efficiency. For each genomic window, read depths were mean-scaled by dividing each position’s depth by the mean depth across all positions within that window, yielding normalized values centered around 1.0, where values greater than 1.0 indicate above-average coverage and values less than 1.0 indicate below-average coverage relative to the window mean.

### 5.2 Blacklisting of SNVs and indels

An internal background mask was constructed from the training cohort and consisted of 46 healthy-control plasma samples sequenced with the capture panel. The mask served two purposes: to reduce residual germline signal in the plasma-only training data (no matched PBMC sequencing was available) and to identify recurrent technical noise.

A two-stage procedure was applied. First, loci with cAF *>* 10% in any control sample were removed (high-background sites), excluding 38 SNV loci and 26 indel loci in the Roche data and 61 SNV loci and 2 indel loci in the Twist data. Second, among the remaining loci, positions with cAF *>* 1% in more than one control sample were blacklisted (recurrent low-level noise), excluding an additional 77 SNV loci and 26 indel loci in the Roche data and 59 SNV loci and 31 indel loci in the Twist data. Masked loci were excluded from model training. Loci flagged as recurrent noise were additionally stored and removed from mutation catalogues during prediction.

In deep targeted UMI sequencing of PBMC, variants were excluded if coverage was below 1000 or if cAF exceeded 0.1% in the PBMC.

### 5.3 Neural network model

#### 5.3.1 Parameterization and training

The SNV model outputs a four-class distribution over {*A, C, G, T*} as in DREAMS, and the probability assigned to a non-reference base is interpreted as the substitution error rate [6]. The indel model outputs a three-class distribution over {*M, I, D*}, where *M* denotes any base observation at the locus (including matches and SNV mismatches) and *I* and *D* denote insertion and deletion observations, respectively.

For the indel model, let *X*_*ij*_ be the observed class for read *j* at genomic position *i*, and let **d**_*ij*_ be the associated feature vector. Each observation is modelled as a single draw from a categorical distribution over {*M, I, D*}. Given a collection of observations {(*x*_*ij*_, **d**_*ij*_)}, the model maximizes the log-likelihood

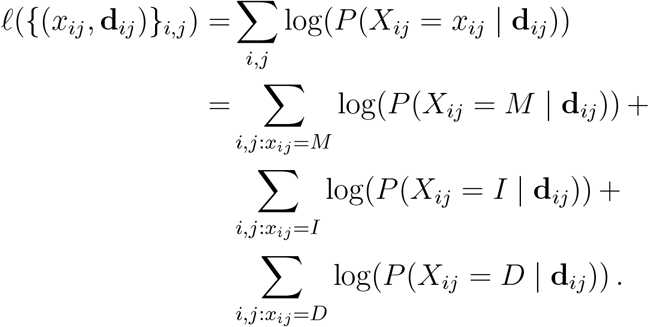

Class probabilities are parameterized with a Softmax over three logits produced by a multilayer perceptron,

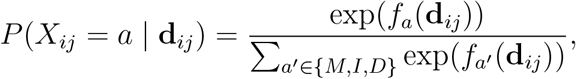

where *f*_*a*_(**d**_*ij*_) is the logit for outcome *a* computed from the final hidden layer. Training minimizes the negative log-likelihood, equivalent to the cross-entropy between observed class labels and the predicted distribution.

The SNV model is defined analogously over four outcomes {*A, C, G, T*} and is trained with the same objective. For downstream inference, predicted base probabilities are collapsed to the binary quantities required by the statistical model.

#### 5.3.2 Architecture

Both the SNV and indel models were implemented as multilayer perceptrons. The input dimension matched the number of selected features. Each network comprised three fully connected hidden layers with 512 units per layer and ReLU activations. Dropout was applied during training with rates of 0.2402 for the SNV model and 0.4658 for the indel model. The output layer was task-specific and parameterized with a Softmax: four outputs for the SNV model corresponding to A, C, G, T, and three outputs for the indel model corresponding to M, I, D. Models were trained using mini-batches of size 2848 (SNV) and 760 (indel) with learning rates of 3.356 ×10^−4^ and 1.736 ×10^−3^, respectively.

The SNV model used the features read index, strand, trinucleotide context, first in pair, UMI count, sequence length, number of other errors, local GC, window ID, and relative position. The indel model used read index, strand, trinucleotide context, sequence length, fragment size, number of other errors, local GC, and window ID.

#### 5.3.3 Feature handling/embedding

Features were partitioned into numerical, categorical, and embedded variables and were preprocessed with a pipeline that was fitted on the training split only and then reused unchanged for validation and test, which ensured consistent input dimensionality and prevented leakage. Numerical features were scaled per feature to match their distribution: approximately Gaussian variables were standard scaled, while count like or heavy tailed variables were transformed with log(1 + *x*) and then robust scaled. A small set of features that were already normalized (for example relative position or residual type quantities) was kept unscaled and only cast to `float32`. Categorical features, including window ID, were one hot encoded using a single encoder that was fitted on the training data and persisted; during validation and test the same encoder was reused with `handle unknown=ignore` to guarantee a fixed output shape even if a split contained only a subset of categories. Finally, the trinucleotide context was handled as an embedded feature as described in [6]. Invalid or missing contexts were mapped to a default index to make preprocessing robust.

### 5.4 Per-mutation score contribution of SNVs and indels

To compare the contribution of SNVs and indels to the DREAMS detection score on a permutation basis, a normalised score was computed for each mutation type separately in the validation cohort. For each CRC patient, PyDREAMS was run twice — once using only SNV loci (alt *∈* {*A, T, C, G*}) and once using only indel loci (alt *∈* {*I, D*}) — to obtain variant-type-specific p-values (*p*_SNV_ and *p*_indel_). The total score per variant type was defined as −log_10_(*p*), and the per-mutation contribution was calculated as −log_10_(*p*)*/n*, where *n* is the number of mutations of the corresponding type in the patient’s tumor-informed panel. Only patients carrying at least one SNV and at least one indel were included in this analysis, so that each patient contributed a paired observation.

### 5.5 Software

The `pydreams` software was implemented in Python and released as an open source package. The source code, documentation, and usage examples were made available on GitHub at https://github.com/mathildediekema/PyDREAMS. Dependency management and reproducible environments were handled with Poetry (`pyproject.toml`), with the package distributed under the name `pydreams` (version 0.1.0) and tested with Python 3.12. Core dependencies included PyTorch for model implementation and training, NumPy and pandas for data handling, scikit-learn for preprocessing (scalers and one hot encoding), and domain specific tooling such as `pysam` for sequencing related workflows. Exact dependency versions were recorded through Poetry to support reproducible installation across systems.

### 5.6 Data Availability

The data underlying this study contain sensitive patient information and are not publicly available due to ethical and confidentiality constraints. Access to data may be available from the corresponding author upon reasonable request and subject to institutional approval and patient consent restrictions.

## Acknowledgment

This work was supported by Danish Data Science Academy, which is funded by the Novo Nordisk Foundation (NNF21SA0069429) and VILLUM FONDEN (40516). This work was also funded by Sygesikring Danmark (2020-0412).

## Author Contributions

M.H.D. wrote the manuscript and performed all analyses and results interpretation. S.O.D. developed the original DREAMS method and provided statistical guidance at the project initiation. C.L.A. leads the colorectal cancer research group, provided patient data and clinical guidance, and reviewed the manuscript. M.H.R. and A.F. generated the sequencing data and guided the experimental process. J.S.P. supervised the study, guided the scientific direction throughout the project, and reviewed and revised the manuscript. All authors reviewed and approved the final manuscript.

## A Appendix

### A.1 Neural network

**Figure A1.**
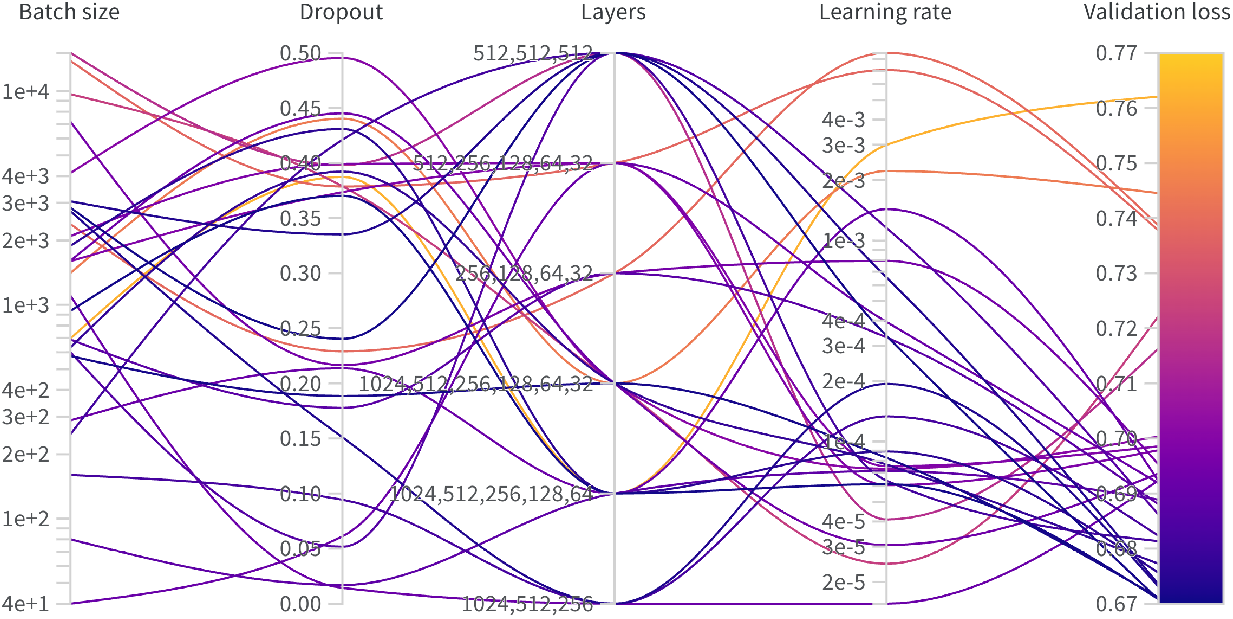
SNV hyperparameter tuning.

**Figure A2.**
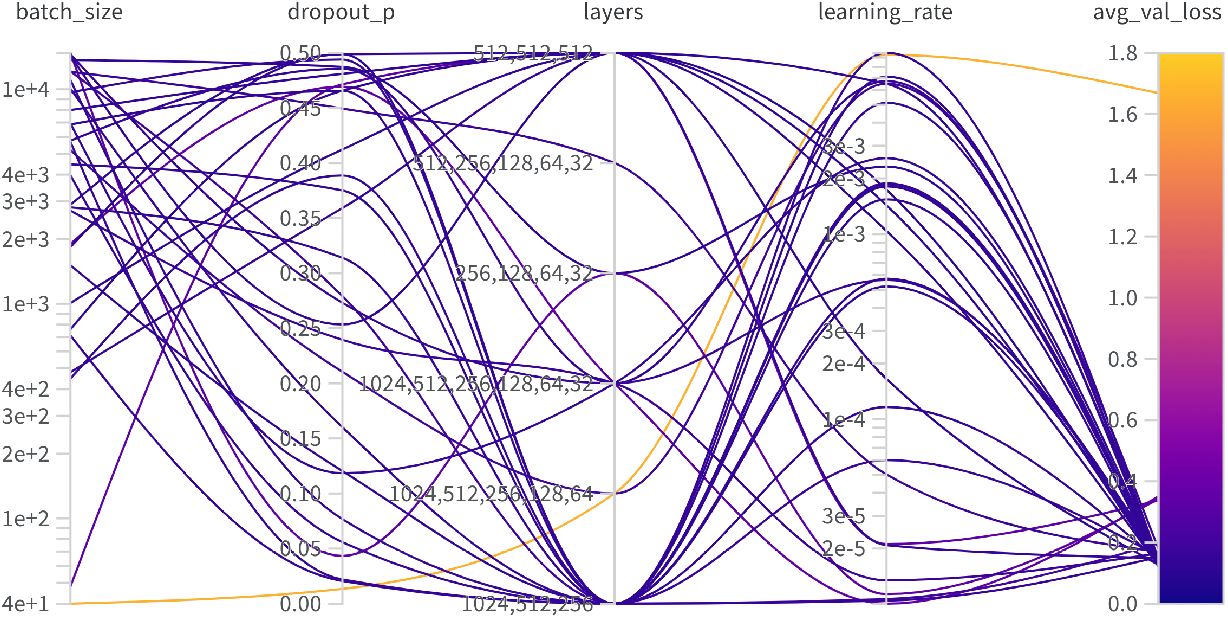
INDEL hyperparameter tuning.

**Table A1:** Neural network hyperparameters for the SNV and indel error models.

| Hyperparameter | SNV model | Indel model |
| --- | --- | --- |
| Dropout probability | 0.2402 | 0.4658 |
| Batch size | 2848 | 760 |
| Learning rate | $3 \times 10^{-4}$ | $2 \times 10^{-3}$ |
| Hidden layers | [512, 512, 512] | [512, 512, 512] |

**Figure A3.**
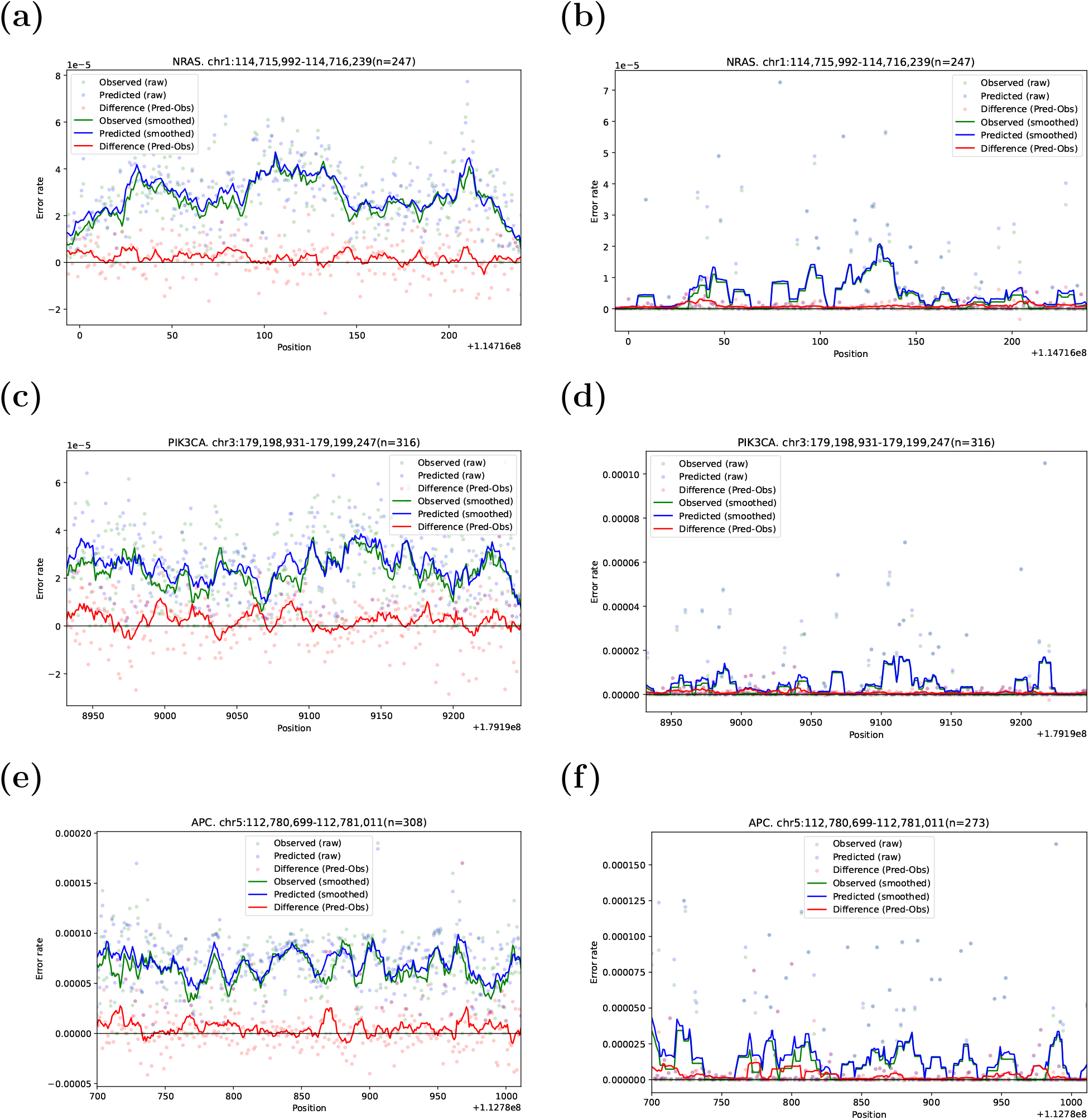
Observed vs predicted error rates in different genomic regions.

#### A.2 Evaluation in test cohort

**Figure A4.**
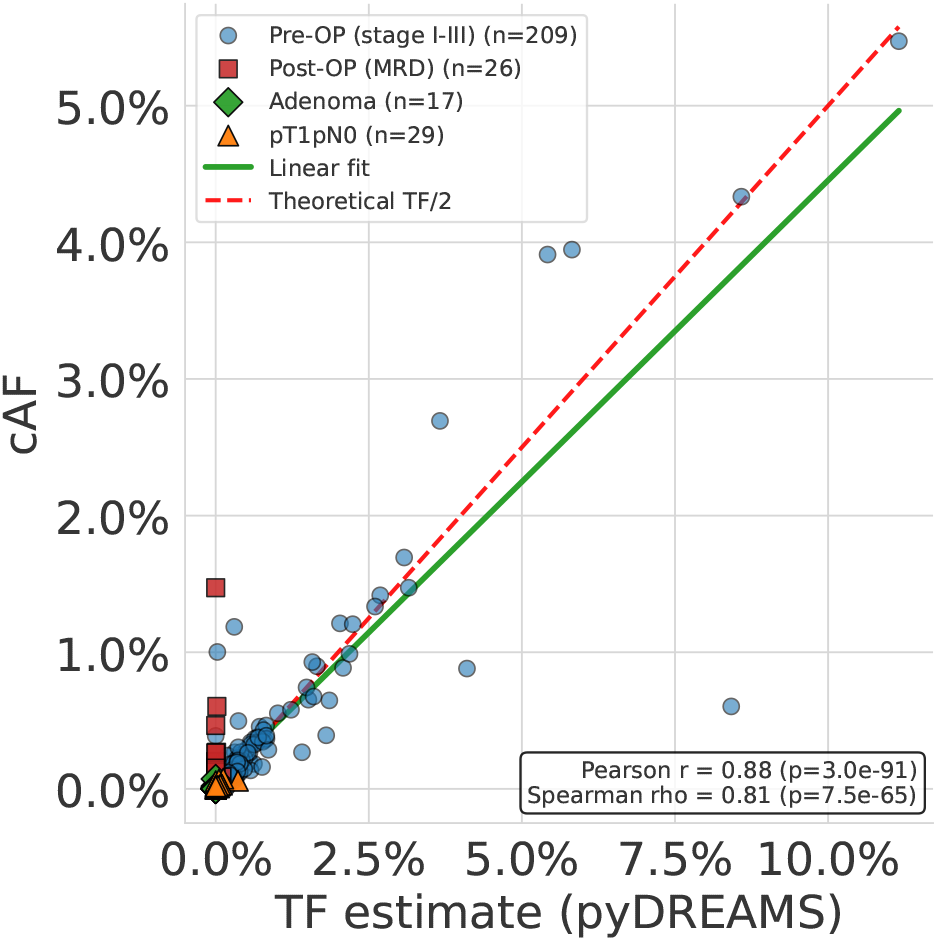
PyDREAMS tumor estimate.

## References

[1] Janine Aucamp, Abel J Bronkhorst, Christoffel PS Badenhorst, and Piet J Pretorius. The diverse origins of circulating cell-free dna in the human body: a critical re-evaluation of the literature. Biological Reviews, 93(3):1649–1683, 2018.

[2] Lukas Biewald. Experiment tracking with weights and biases, 2020. Software available from wandb.com.

[3] Feifei Cheng, Li Su, and Cheng Qian. Circulating tumor dna: a promising biomarker in the liquid biopsy of cancer. Oncotarget, 7(30):48832, 2016.

[4] Xuanjin Cheng, Murathan T Goktas, Laura M Williamson, Martin Krzywinski, David T Mulder, Lucas Swanson, Jill Slind, Jelena Sihvonen, Cynthia R Chow, Amy Carr, et al. Enhancing clinical genomic accuracy with panelgc: a novel metric and tool for quantifying and monitoring gc biases in hybridization capture panel sequencing. Briefings in Bioinformatics, 25(5), 2024.

[5] Re-I Chin, Kevin Chen, Abul Usmani, Chanelle Chua, Peter K Harris, Michael S Binkley, Tej D Azad, Jonathan C Dudley, and Aadel A Chaudhuri. Detection of solid tumor molecular residual disease (mrd) using circulating tumor dna (ctdna). Molecular diagnosis & therapy, 23(3):311–331, 2019.

[6] Mikkel H Christensen, Simon O Drue, Mads H Rasmussen, Amanda Frydendahl, Iben Lyskjær, Christina Demuth, Jesper Nors, Kåre A Gotschalck, Lene H Iversen, Claus L Andersen, et al. Dreams: deep read-level error model for sequencing data applied to low-frequency variant calling and circulating tumor dna detection. Genome Biology, 24(1):99, 2023.

[7] Han Fang, Yiyang Wu, Giuseppe Narzisi, Jason A ORawe, Laura T Jimenez Barrón, Julie Rosenbaum, Michael Ronemus, Ivan Iossifov, Michael C Schatz, and Gholson J Lyon. Reducing indel calling errors in whole genome and exome sequencing data. Genome medicine, 6(10):89, 2014.

[8] Aaron Fisher, Cynthia Rudin, and Francesca Dominici. All models are wrong, but many are useful: Learning a variable’s importance by studying an entire class of prediction models simultaneously. Journal of Machine Learning Research, 20(177):1–81, 2019.

[9] Tim Forshew, Muhammed Murtaza, Christine Parkinson, Davina Gale, Dana WY Tsui, Fiona Kaper, Sarah-Jane Dawson, Anna M Piskorz, Mercedes Jimenez-Linan, David Bentley, et al. Noninvasive identification and monitoring of cancer mutations by targeted deep sequencing of plasma dna. Science translational medicine, 4(136):136ra68–136ra68, 2012.

[10] Amanda Frydendahl, Mads Heilskov Rasmussen, Sarah Østrup Jensen, Tenna Vesterman Henriksen, Christina Demuth, Mathilde Diekema, Henrik Jørn Ditzel, Sara Witting Christensen Wen, Jakob Skou Pedersen, Lars Dyrskjøt, et al. Error-corrected deep targeted sequencing of circulating cell-free dna from colorectal cancer patients for sensitive detection of circulating tumor dna. International Journal of Molecular Sciences, 25(8):4252, 2024.

[11] Ellen Heitzer, Imran S Haque, Charles ES Roberts, and Michael R Speicher. Current and future perspectives of liquid biopsies in genomics-driven oncology. Nature Reviews Genetics, 20(2):71–88, 2019.

[12] Isaac Kinde, Jian Wu, Nick Papadopoulos, Kenneth W Kinzler, and Bert Vogelstein. Detection and quantification of rare mutations with massively parallel sequencing. Proceedings of the National Academy of Sciences, 108(23):9530–9535, 2011.

[13] Anatoli Kustanovich, Ruth Schwartz, Tamar Peretz, and Albert Grinshpun. Life and death of circulating cell-free dna. Cancer biology & therapy, 20(8):1057–1067, 2019.

[14] Jing Lei, Max G’Sell, Alessandro Rinaldo, Ryan J Tibshirani, and Larry Wasserman. Distribution-free predictive inference for regression. Journal of the American Statistical Association, 113(523):1094–1111, 2018.

[15] Shuo Li, Weihua Zeng, Xiaohui Ni, Yonggang Zhou, Mary L Stackpole, Zorawar S Noor, Zuyang Yuan, Adam Neal, Sanaz Memarzadeh, Edward B Garon, et al. cftrack: a method of exome-wide mutation analysis of cell-free dna to simultaneously monitor the full spectrum of cancer treatment outcomes including mrd, recurrence, and evolution. Clinical Cancer Research, 28(9):1841–1853, 2022.

[16] Christoffer Trier Maansson, Anders Lade Nielsen, and Boe Sandahl Sorensen. Liquid biopsy epigenetics: establishing a molecular profile based on cell-free dna. Molecular Oncology, 2025.

[17] Tina Moser, Stefan Kuehberger, Isaac Lazzeri, Georgios Vlachos, and Ellen Heitzer. Bridging biological cfdna features and machine learning approaches. Trends in Genetics, 39(4):285–307, 2023.

[18] Giuseppe Narzisi and Michael C Schatz. The challenge of small-scale repeats for indel discovery. Frontiers in bioengineering and biotechnology, 3:8, 2015.

[19] Aaron M Newman, Scott V Bratman, Jacqueline To, Jacob F Wynne, Neville CW Eclov, Leslie A Modlin, Chih Long Liu, Joel W Neal, Heather A Wakelee, Robert E Merritt, et al. An ultrasensitive method for quantitating circulating tumor dna with broad patient coverage. Nature medicine, 20(5):548–554, 2014.

[20] Aaron M Newman, Alexander F Lovejoy, Daniel M Klass, David M Kurtz, Jacob J Chabon, Florian Scherer, Henning Stehr, Chih Long Liu, Scott V Bratman, Carmen Say, et al. Integrated digital error suppression for improved detection of circulating tumor dna. Nature biotechnology, 34(5):547–555, 2016.

[21] David A Nix, Sabine Hellwig, Christopher Conley, Alun Thomas, Carrie L Fuertes, Cindy L Hamil, Preetida J Bhetariya, Ignacio Garrido-Laguna, Gabor T Marth, Mary P Bronner, et al. The stochastic nature of errors in next-generation sequencing of circulating cell-free dna. PLoS One, 15(2):e0229063, 2020.

[22] Nathan D Olson, Justin Wagner, Nathan Dwarshuis, Karen H Miga, Fritz J Sedlazeck, Marc Salit, and Justin M Zook. Variant calling and benchmarking in an era of complete human genome sequences. Nature Reviews Genetics, 24(7):464–483, 2023.

[23] Luciana Santos Pessoa, Manoela Heringer, and Valéria Pereira Ferrer. ctdna as a cancer biomarker: A broad overview. Critical reviews in oncology/hematology, 155:103109, 2020.

[24] Satyanarayan Rao, Amy L Han, Alexis Zukowski, Etana Kopin, Carol A Sartorius, Peter Kabos, and Srinivas Ramachandran. Transcription factor–nucleosome dynamics from plasma cfdna identifies er-driven states in breast cancer. Science Advances, 8(34):eabm4358, 2022.

[25] Jesse J Salk, Michael W Schmitt, and Lawrence A Loeb. Enhancing the accuracy of next-generation sequencing for detecting rare and subclonal mutations. Nature Reviews Genetics, 19(5):269–285, 2018.

[26] Cynthia Sanchez, Matthew W Snyder, Rita Tanos, Jay Shendure, and Alain R Thierry. New insights into structural features and optimal detection of circulating tumor dna determined by single-strand dna analysis. NPJ genomic medicine, 3(1):31, 2018.

[27] Seung-Ho Shin, Woong-Yang Park, and Donghyun Park. Characterization of dna lesions associated with cell-free dna by targeted deep sequencing. BMC Medical Genomics, 14(1):192, 2021.

[28] Matthew W Snyder, Martin Kircher, Andrew J Hill, Riza M Daza, and Jay Shendure. Cell-free dna comprises an in vivo nucleosome footprint that informs its tissues-of-origin. Cell, 164(1):57–68, 2016.

[29] Nicholas Stoler and Anton Nekrutenko. Sequencing error profiles of illumina sequencing instruments. NAR genomics and bioinformatics, 3(1):lqab019, 2021.

[30] Ge Tan, Lennart Opitz, Ralph Schlapbach, and Hubert Rehrauer. Long fragments achieve lower base quality in illumina paired-end sequencing. Scientific reports, 9(1):2856, 2019.

[31] Todd J Treangen and Steven L Salzberg. Repetitive dna and next-generation sequencing: computational challenges and solutions. Nature Reviews Genetics, 13(1):36–46, 2012.

[32] Peter Ulz, Samantha Perakis, Qing Zhou, Tina Moser, Jelena Belic, Isaac Lazzeri, Albert Wölfler, Armin Zebisch, Armin Gerger, Gunda Pristauz, et al. Inference of transcription factor binding from cell-free dna enables tumor subtype prediction and early detection. Nature communications, 10(1):4666, 2019.

[33] Jonathan CM Wan, Charles Massie, Javier Garcia-Corbacho, Florent Mouliere, James D Brenton, Carlos Caldas, Simon Pacey, Richard Baird, and Nitzan Rosenfeld. Liquid biopsies come of age: towards implementation of circulating tumour dna. Nature Reviews Cancer, 17(4):223–238, 2017.

[34] Adam J Widman, Minita Shah, Amanda Frydendahl, Daniel Halmos, Cole C Khamnei, Nadia Øgaard, Srinivas Rajagopalan, Anushri Arora, Aditya Deshpande, William F Hooper, et al. Ultrasensitive plasma-based monitoring of tumor burden using machine-learning-guided signal enrichment. Nature medicine, 30(6):1655–1666, 2024.

[35] Guanhua Zhu, Chowdhury Rafeed Rahman, Victor Getty, Denis Odinokov, Probhonjon Baruah, Hanaé Carrié, Avril Joy Lim, Yu Amanda Guo, Zhong Wee Poh, Ngak Leng Sim, et al. A deep-learning model for quantifying circulating tumour dna from the density distribution of dna-fragment lengths. Nature Biomedical Engineering, pages 1–13, 2025.

